# Combining multiple functional annotation tools increases coverage of metabolic annotation

**DOI:** 10.1101/160887

**Authors:** Marc Griesemer, Jeffrey Kimbrel, Carol Zhou, Ali Navid, Patrik D’haeseleer

**Author notes:** The authors wish it to be known that, in their opinion, the first 2 authors should be regarded as joint First Authors.

## Abstract

Genome-scale metabolic modeling is a cornerstone of systems biology analysis of microbial organisms and communities, yet these genome-scale modeling efforts are invariably based on incomplete functional annotations. Annotated genomes typically contain 30-50% of genes without functional annotation, severely limiting our knowledge of the “parts lists” that the organisms have at their disposal. These incomplete annotations may be sufficient to derive a model of a core set of well-studied metabolic pathways that support growth in pure culture. However, pathways important for growth on unusual metabolites exchanged in complex microbial communities are often less understood, resulting in missing functional annotations in newly sequenced genomes. Here, we present results on a comprehensive reannotation of 27 bacterial reference genomes, focusing on enzymes with EC numbers annotated by KEGG, RAST, EFICAz, and the BRENDA enzyme database, and on membrane transport annotations by TransportDB, KEGG and RAST. Our analysis shows that annotation using multiple tools can result in a drastically larger metabolic network reconstruction, adding on average 40% more EC numbers, 3-8 times more substrate-specific transporters, and 37% more metabolic genes. These results are even more pronounced for bacterial species that are more phylogenetically distant from well-studied model organisms such as E. coli.

## INTRODUCTION

In the early days of genome sequencing, functional annotation involved computational prediction of gene function coupled with extensive manual curation by teams of experts (1–3). Today, with the exponential explosion of DNA sequencing (4) the fraction of genes that have undergone any degree of manual curation or even experimental validation is becoming vanishingly small (5, 6). Automated gene annotation tools employing different methodologies with minimal manual curation are widely used, functionally annotating by homology to existing annotations, or by identification of conserved domains/motifs within a coding sequence (7, 8). Many draft genomes and metagenome bins are often run through a single annotation pipeline where genome annotations are inherited from previous genome annotations. Even when multiple annotation tools are used, integrating the different outputs in a cohesive manner remains a major challenge (9, 10). However, individual metabolic annotation tools often return annotations for different subsets of genes, offering the potential to greatly increase the coverage of metabolic annotations by combining the outputs of multiple tools if the barrier for integration can be overcome.

“Genome-scale” metabolic models implicitly assume complete and accurate functional annotation. However, 30-50% of genes in a typical genome still lack any functional annotation (11), a statistic which has not improved much over the past two decades of genome sequencing (3). More than 30% of these unannotated genes are estimated to have metabolic functions (12) leaving a significant gap in our understanding of the underlying metabolic processes. In addition, annotated genes include a large (and potentially growing) fraction of misannotations (13). Further, high-throughput untargeted metabolomics often contain a large fraction of peaks that cannot be reliably matched to any known metabolites (14), and many of those that can be identified often do not match any metabolic reconstruction of the microbial species involved (15), providing another strong indication of the extent of microbial metabolism we are missing with traditional metabolic annotation methods. Metabolic modeling efforts, however, are moving beyond studying core metabolic pathways in a single organism towards multi-species models, real-world communities and ecosystems (16–18), and incorporation of complex ‘omics and metabolite data, emphasizing the need for a more complete coverage of the metabolic functions identified in microbial genomes.

In constraint-based genome-scale modeling methods such as Flux Balance Analysis (19), the issue of missing metabolic annotations is dealt with by “gap filling” - the addition of a set of metabolic reactions beyond those that were derived directly from the genome annotation (20). A variety of gap filling algorithms have been developed to predict the missing reactions necessary to make the metabolic network model sufficiently complete to produce biomass (21–24). In a broad collection of 130 genome-scale metabolic models added to the ModelSEED database (25), on average 56 additional gap filled reactions were needed for each model to produce biomass in simple defined nutrient media. Even after those additions an average of one-third of the reactions in each model were still blocked, meaning that there were still enough reactions missing in the network to preclude metabolic flux through those reactions (25, 26). In addition, the number of reactions that can partake in a gap filling solution is vast (3,270 in the case of *E. coli*), and the sets of reactions generated by different gap filling algorithms may have little or no overlap with each other (27). Clearly, a more complete identification and annotation of metabolic reactions would be preferable to the addition of dozens of poorly supported reactions just to patch the holes in the network.

Recent genome-scale modeling of *Clostridium beijerinckii* NCIMB 8052 (28) demonstrated that the total number of genes and reactions included in the final curated model could be almost doubled by incorporating multiple annotation tools. The reconstruction of the *C. beijerinckii* metabolic network used 3 different database sources (SEED (29), KEGG (30), and RefSeq annotations captured in BioCyc (31)) to evaluate annotation coverage and produce a more robust model. Each annotation source contributed only about half of the reactions in the final curated model. Only a third of the reactions were present in all three sources, and these reactions were found to contribute significantly to a core set of active reactions in validation simulations. The small overlap between annotations was not simply due to any source contributing more heavily to a particular area of metabolism, nor did any one source outperform another in terms of model connectivity. Likewise, in an analysis of nine prokaryotic genomes, the three enzyme annotation sources used – NCBI, KEGG, and the PEDANT protein database (32) – only agreed on less than one third of the annotated genes (33).

Accurate metabolic models also rely on accurate determination of substrate transport between the bacterium and its environment. Transporter annotations have rarely been used in genome scale metabolic modeling because of the difficulty in computationally determining the exact substrate being transported. Because of this, many metabolic modeling methods simply assume that a transporter exists for the import of any necessary metabolite - an assumption that is incorrect in some cases. For example, the yogurt bacteria *Streptococcus thermophilus* has an unusual growth phenotype in that it grows poorly on glucose, even though it possesses all the required metabolic enzymes (34). Instead it preferably imports the disaccharide lactose, hydrolyzes it to glucose and galactose, then secretes the galactose back out of the cell. S. *thermophilus* lacks the typical glucose phosphotransferase system used by many bacteria, and instead has an efficient lactose import mechanism that makes it well adapted to grow in milk (34). Prediction tools such as TransportDB’s Transporter Automatic Annotation Pipeline (TransAAP, (35)) now allow researchers to generate substrate predictions that are sufficiently detailed to be included in metabolic pathways, and could give insights into growth or metabolite exchange phenotypes that are not readily apparent from the metabolic pathways present in the genome.

We undertook an investigation into the effectiveness of several popular tools for genome annotation and their overlap with each other, focusing specifically on enzymatic annotations characterized by EC (Enzyme Commission) numbers, because those can be most unambiguously mapped across annotations from different sources (36). Note that many databases such as RAST, KEGG and MetaCyc also curate their own set of metabolic reactions and reaction variants beyond the canonical EC number hierarchy, however, differing reaction identifiers can be much more difficult to compare across the different tools. Using 27 bacterial reference genomes from BioCyc (31), we evaluated how many genes, EC numbers, and gene-EC annotations were unique or shared with other tools. We also undertook a study of how transporter annotations were handled between RAST (29), KEGG (30), and TransportDB (35), focusing especially on transporters with detailed substrate annotations. We hypothesized that by combining annotation tools we could alleviate some of the known problems with lack of coverage of metabolic annotations, especially for less well studied organisms and pathways, and transporters.

## MATERIAL AND METHODS

### Reference Genomes

We focused on a total of 27 genomes of BioCyc Tier 1 & Tier 2 bacteria (31). These genomes are from a range of phyla, including 15 Proteobacteria, 6 Firmicutes, 3 Actinobacteria, 2 Bacteroidetes and 1 Cyanobacteria. In addition to a range of phyla, these 27 organisms also cover different lifestyles, including a human gut symbiont (*Bacteroides thetaiotaomicron* VPI-5482), pathogens (e.g. *Mycobacterium tuberculosis*), an obligate insect endosymbiont (*Candidatus* Evansia muelleri), an unusual manganese oxidizing bacterium (*Aurantimonas manganoxydans* SI85-9A1), and a photoautotrophic cyanobacterium (*Synechococcus elongatus* PCC 7942). Genbank files were downloaded from NCBI (accessions listed in Table 1), and standardized to remove all functional annotation, retaining only the original open reading frames and locus tag/protein identification information (Supplementary Data file S3).

**Table 1.**
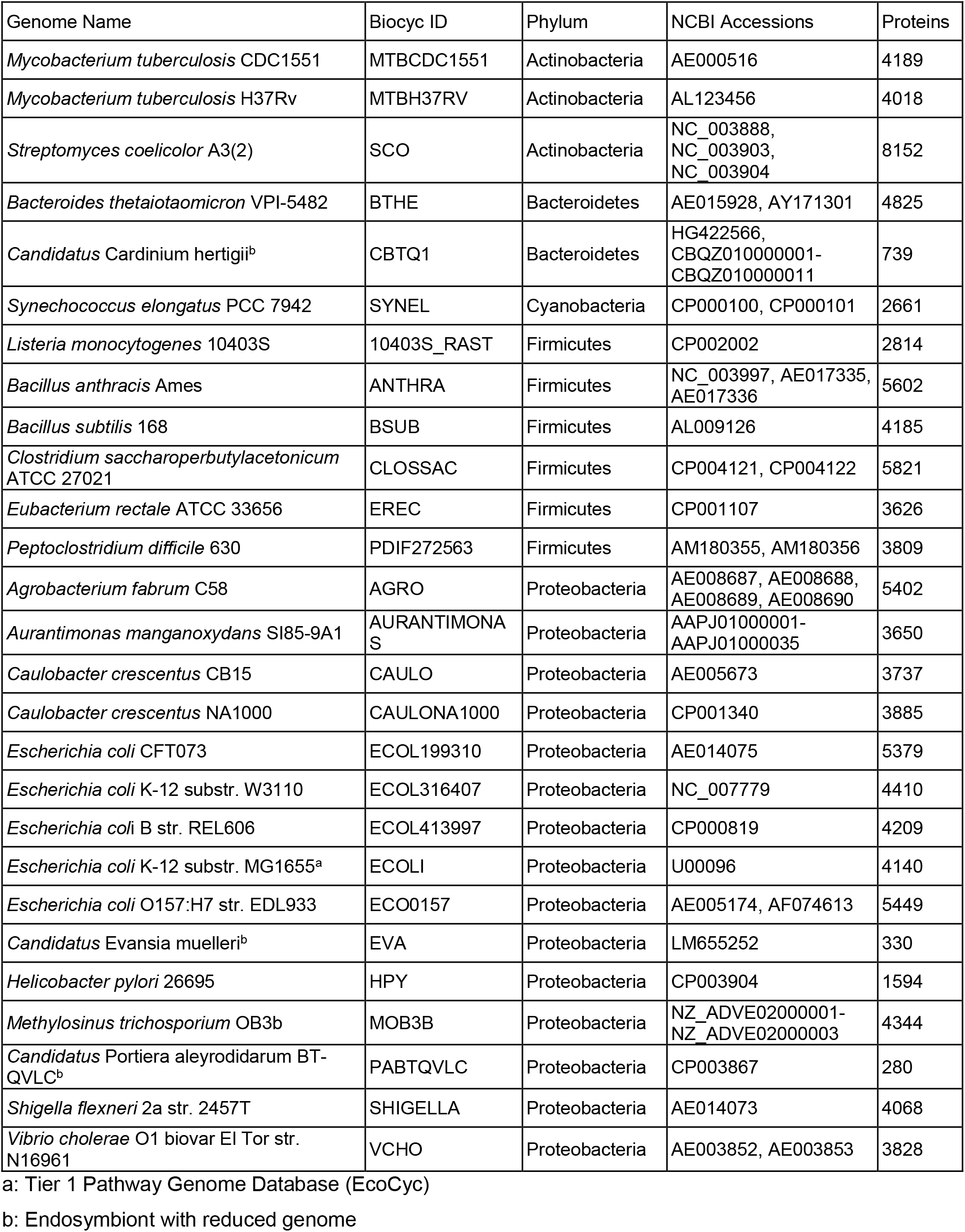
Reference genomes used in this study

### Annotation Tools

#### RAST

(Rapid Annotation Subsystem Technology, (29)) is an open-source web server for genome annotation, using an assignment propagation strategy based on manually curated subsystems and subsystem-based protein families that automatically guarantees a high degree of assignment consistency. RAST returns an analysis of the genes and subsystems in each genome. We used the NMPDR website (rast.nmpdr.org) to generate genome-wide annotations for our 27 reference genomes and parsed any EC numbers from the functional annotation. This is also the core metabolic annotation tool used by the popular ModelSEED tool for generating draft genome-scale models of metabolism (37).

#### KEGG

(Kyoto Encyclopedia of Genes and Genomes, (30)) is a collection of genome and pathway databases for systems biology. We used KAAS (KEGG Automatic Annotation Server, http://www.genome.jp/tools/kaas, (38)) to generate genome-wide annotations for our 27 reference genomes. KAAS assigns KEGG Orthology (KO) numbers using the bi-directional best hit method (BBH) against a set of default prokaryotic genomes in the KEGG database. We mapped KO numbers to EC numbers using a mapping table provided by the KEGG BRITE Database (http://www.genome.jp/kegg-bin/get_htext?ko01000.keg).

#### EFICAz

(39) uses large scale inference to classify enzymes into functional families, combining 4 methods into a single approach without the need for structural information. Recognition of functionally discriminating residues (FDR) allows EFICAz to use a method called evolutionary footprinting. EFICAz has been rigorously crossed validated, and achieves very high precision and recall, even for sequence similarities to known enzymes as low as 40%. We used a local install of EFICAz^2.5^ to generate EC number predictions for our 27 reference genomes.

#### BRENDA

(40) is a database containing 7.2 million enzyme sequences categorized into 82,568 enzymatic functions based on the literature and contains functional and molecular information such as nomenclature, structure, and substrate specificity. We annotated our 27 reference genomes based on a BLAST search at >60% sequence identity against a local copy of the 2011 BRENDA database of enzyme reference sequences.

All EC number annotations are available in Supplementary Data file S4.

### Transport Annotations

Where available, pre-generated transporter annotations were downloaded from the TransportDB 2.0 database (35). For those reference genomes that were not already present in the database (*M. trichosporium* OB3b, *Candidatus* C. hertigii, *Candidatus* E. muelleri, and *A. manganoxydans* SI85-9A1), we submitted the predicted protein sequences to the TransAAP web-based transporter annotation tool using the default parameters (http://www.membranetransport.org/transportDB2/TransAAP_login.html). We also retrieved transporter annotations from RAST and KEGG. For RAST, annotations were filtered for the subsystem for “Membrane Transport”. For KEGG, we mapped KO numbers to transporter annotations using a mapping table provided by the KEGG BRITE Database (http://www.genome.jp/kegg-bin/get_htext?ko02000.keg). Substrate names were ranked from most to least specific (see Table 2). Substrates that can be incorporated as a transport reaction in a metabolic model were ranked 1 and 2. Broader substrate classes that could be used for gap filling or interpretation of transcriptomics data were ranked 3 and 4, and annotated transporters without substrate prediction were ranked 5. The full table of substrate names and ranking can be found in Supplementary Data file S6.

**Table 2.**
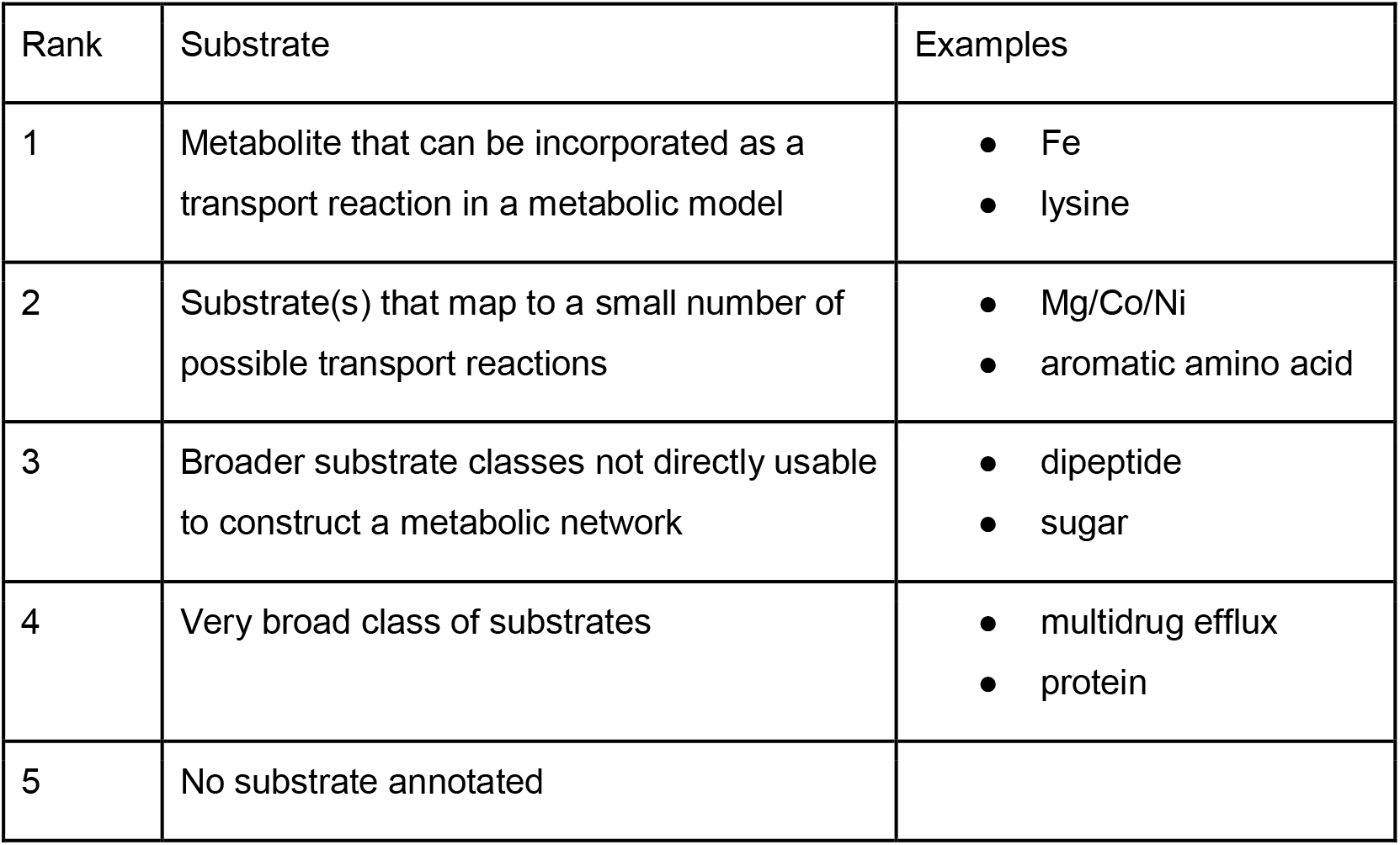
Examples of substrate annotation ranking, from most specific (rank 1) to least specific (no substrate, rank 5). See Supplementary Data file S7 for the full table.

## RESULTS AND DISCUSSION

### Discrepancies in metabolic annotations between different tools

In total, the RAST, KAAS, EFICAz and BRENDA tools produced 47,447 Gene-EC annotations (“gene X codes for an enzyme with EC number Y”) across the 27 reference genomes, for an average of 1,757 annotations per genome. The metabolic gene-EC annotations produced by these automated genome-wide annotation tools differed drastically (Figure 1). Each tool produced on average between 23% (EFICAz) and 48% (BRENDA) unique gene-EC annotations that were not predicted by any of the other tools. Overall, fewer than a quarter of all gene-EC annotations were agreed on by at least 3 tools.

**Figure 1.**
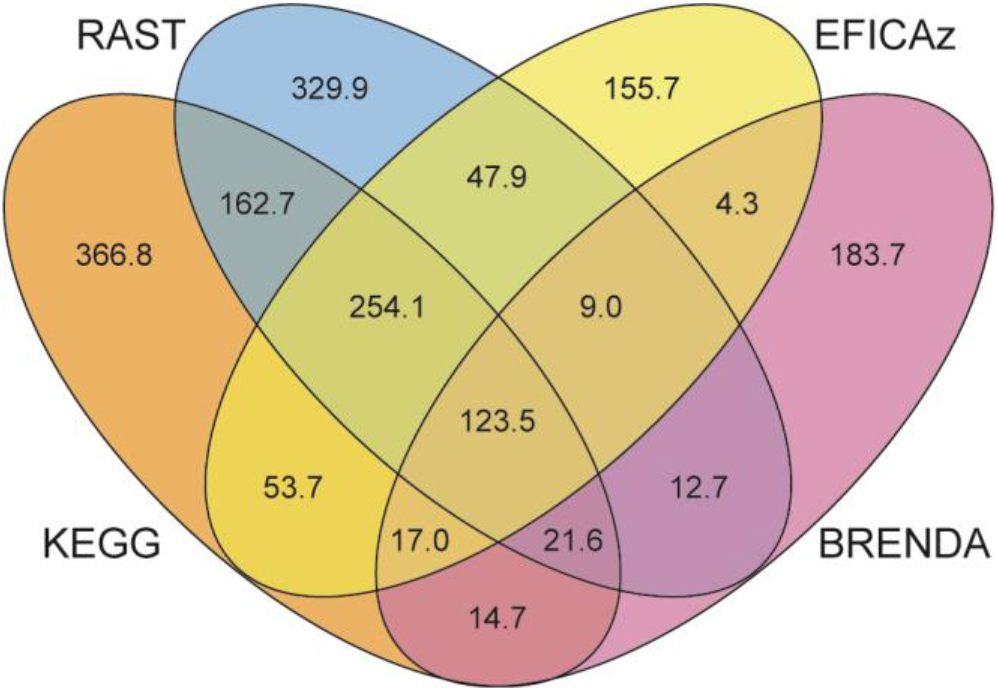
Large differences exist between the sets of Gene-EC annotations generated by the four annotation tools across the 27 reference genomes.

When two annotation tools both assigned a particular gene an EC annotation, the two tools generally agreed and assigned at least one identical EC annotation in more than 50% of cases (Table 3). BRENDA on average had the lowest agreement with other tools (56.0%-69.7%). Note also that BRENDA had a larger fraction (47.5%) of gene-EC annotations not shared by any other tools (Figure 1). In contrast, EFiCAz showed the highest agreement with other tools (69.7%-86.4%) and had the lowest number of gene-EC annotations not shared by other tools (23.4%).

**Table 3.**
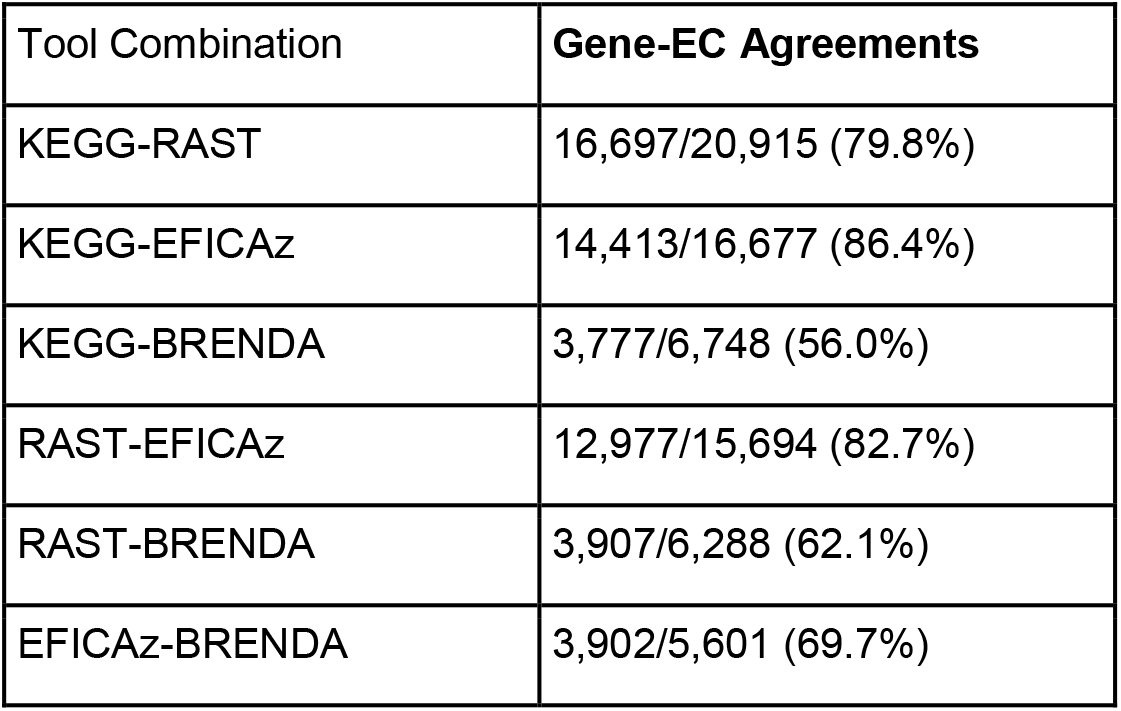
Percentage of gene-EC annotation agreements that exist between pairs of tools. The denominator is the number of genes across the 27 reference genomes that are covered by both tools. The numerator counts the number of such genes for which both tools provide at least one identical gene-EC number annotation.

Comparing annotation tools against each other can give a sense of which tools are closest to a consensus annotation, or which tools seem to be outliers, however assessing the integrity of these predictions is difficult without experimental validation. Therefore, to determine which tool provides the best ratio of true/false annotation predictions we compared their predictions to the EcoCyc database (41). EcoCyc is a gold-standard continuously updated database of experimentally determined and extensively hand-curated enzymatic functions in *Escherichia coli* K-12 substr. MG1655, the most-studied model organism in modern biology. We used the gene-EC numbers annotated in EcoCyc as a set of true positives to evaluate how well the two most commonly used automated annotation tools, RAST and KEGG, are able to assign function to the enzymes in *E. coli* K-12. Overall, there was a high degree of overlap between the RAST and KEGG predictions (Figure 2) with EcoCyc, however neither tool covered all of EcoCyc, and both tools predicted a small number of reactions not experimentally validated.

**Figure 2.**
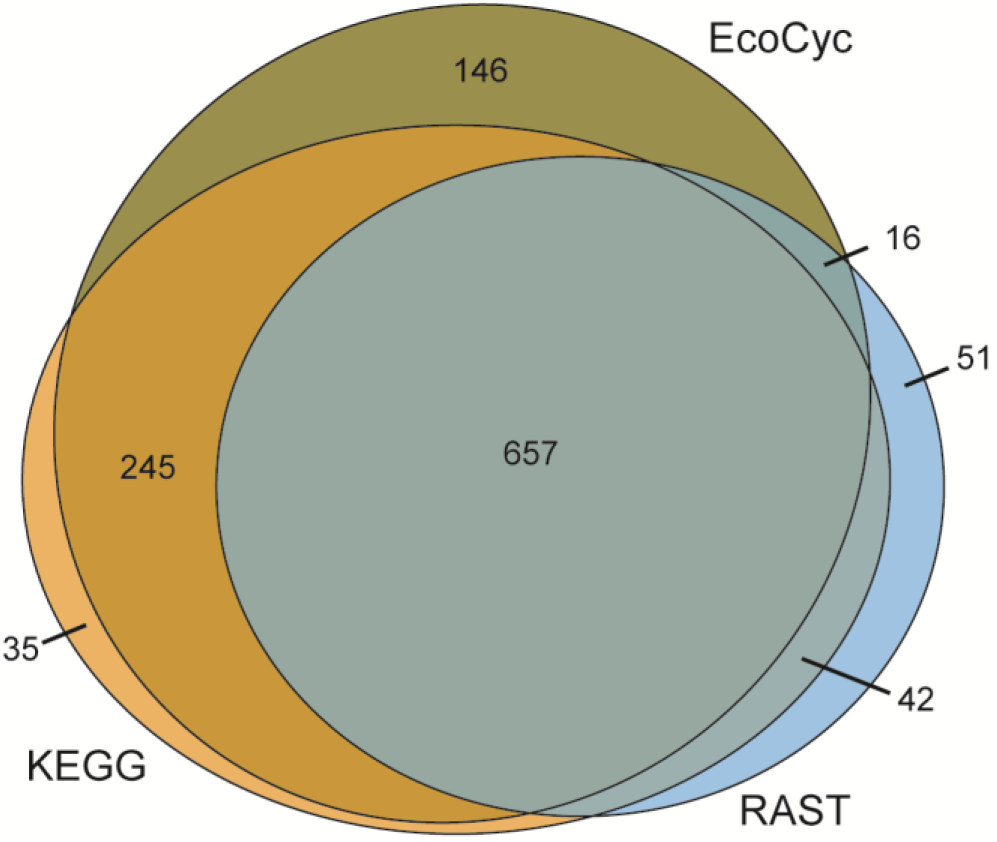
Gene-EC annotations produced by KEGG and RAST for E. coli K-12, compared to the EcoCyc gold standard. The sets and intersections are drawn proportionally to the number of annotations in each.

One major caveat of using *E. coli*to evaluate the quality of annotation tools is that so much of our knowledge of microbial metabolism is based on *E. coli*, and therefore annotation tools can be expected to be trained or optimized on *E. coli* to some extent, so performance on *E. coli* is not necessarily indicative of results on other organisms. For example, the KEGG annotation provided by KAAS is done by calculating bidirectional best BLAST hits against annotated reference genomes including *E. coli*, essentially providing a direct lookup of *E. coli* annotations in the KEGG database.

### Coverage of the metabolic network reconstruction

While the individual gene-EC annotations examined in the previous section reflected the quality and agreement between annotation tools, the total set of EC reactions annotated for a genome by each tool reflects the size and coverage of its metabolic network reconstruction. In this case, we simply counted the total number of different EC numbers, regardless of whether multiple genes are annotated with the same EC number (isozymes), or whether genes were annotated with multiple EC numbers (multifunctional enzymes). On average, the four tools combined produced 868 EC reactions per genome, with the largest agreement between RAST and KEGG (Figure 3). In general, KEGG produced a larger number of unique EC numbers, which could indicate more over-prediction, or more comprehensive pathway coverage. Note that both RAST and KEGG also generate many reactions without official EC numbers, so in some cases these annotation tools may produce annotations that are minor variants or subsets of the canonical EC number reaction in EcoCyc.

**Figure 3.**
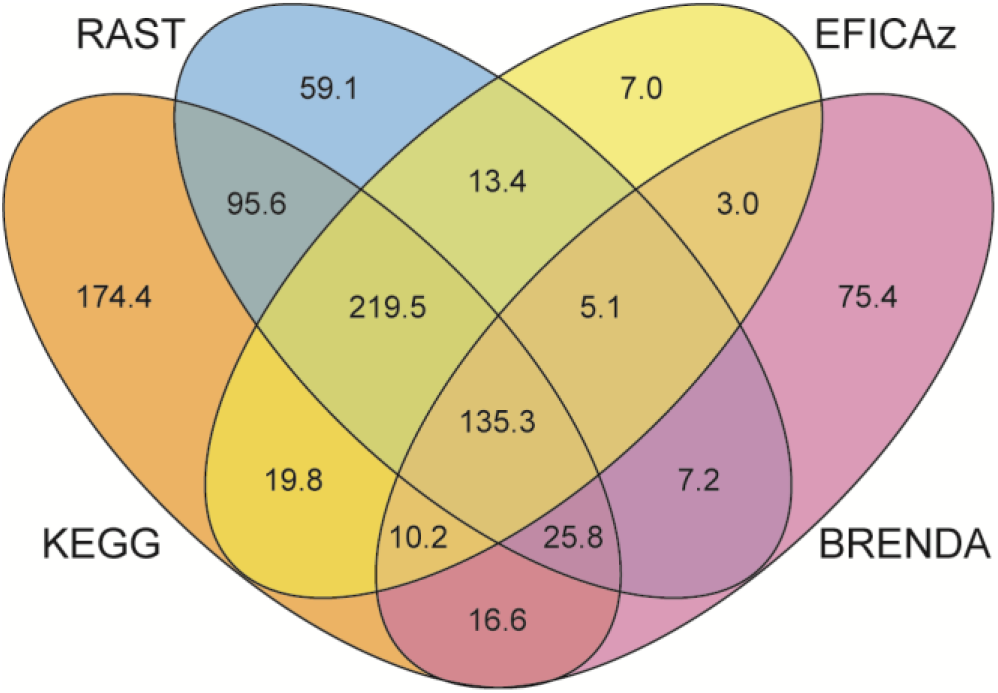
Reaction overlap between the annotation tools (average number of EC numbers per genome).

EFICAz produced the least number of unique EC numbers, but high agreement with RAST and KEGG, suggesting that it can be used in combination with them to highlight high confidence annotations. Note that EFICAz also produces incomplete “three-digit” EC number annotations (*e.g*. 1.2.3.-) which may be useful for hole filling, but were not considered in this analysis.

BLASTing against the BRENDA database of reference enzymes produced the smallest number of annotations, but a high fraction of unique EC numbers. Interestingly, of the top 10 unique EC numbers produced by this method, only one is also covered by RAST and KEGG, two of the EC numbers have been deprecated by the Enzyme Commission, and six are EC numbers that have been assigned in 2000 or later and may not have been incorporated into the predictions by the other annotation tools yet. So even though a simple BLAST against a reference database such as BRENDA proves to be one of the less effective means for assigning metabolic functions, it may still have some value to capture recently described enzymes not already covered by the more sophisticated enzyme prediction tools.

While counting the number of EC numbers reflects the size of the metabolic network, counting the numbers of genes that have received any metabolic annotation reflects the genome annotation coverage. Supplementary Figure S1 shows the number of genes in all of the reference genomes annotated with one or more EC numbers by each of the tools. On average, 1,361 genes per genome were assigned a function with at least one tool, and more than 65% of these genes were assigned a metabolic function by more than one tool. The results show that just as each tool adds a significant number of reactions to the metabolic network model, each tool also significantly contributes to the number of genes covered with metabolic annotations.

The EC numbers on which the different tools most often agree across the 27 reference genomes tended to belong to well-studied core metabolic pathways. Out of the 79 EC numbers on which all four tools agreed in at least half of the genomes (Supplementary Data S5), more than three quarters (61/79) were involved in biosynthesis or biodegradation of amino acids, nucleotides, carbohydrates, and cofactors; or in the processing of RNA, DNA and proteins. In contrast, almost none of these EC numbers were involved in biosynthesis or degradation of fatty acids, lipids, aromatics compounds, or secondary metabolites.

The differing sets of annotations produced by each tool can enable a user to trade off confidence for coverage, with higher confidence obtained when accepting only annotations that were agreed upon by multiple tools (the intersection), or higher coverage obtained by using the combined set of annotations from multiple tools (the union). To examine the effect of taking the intersection (higher confidence) or union (higher coverage) of the annotation tools, we compared combinations of the four annotation tools against the *E. coli* K-12 “gold standard” metabolic reactions in EcoCyc (Figure 4). These combinations included the single tool annotations, as well as the union and intersection of all four tools combined in pairs, triplets and quartets. The resulting EC annotations from these combinations were then compared to the 1,064 EC numbers from EcoCyc, and the count of true positives, false positives, and false negatives were calculated for each combination. *True positives* (TP) correspond to EC numbers predicted by the annotation tools and present in EcoCyc. *False positives* (FP) are EC numbers annotated by the annotation tools, but not found in EcoCyc. *False negatives* (FN) are those EC numbers that are in EcoCyc but were not predicted by the annotation tools. *Precision* is defined as TP/(TP+FP), that is, the fraction of predicted EC numbers that are actually found in EcoCyc. Lower precision indicates an overprediction of EC annotations that are not experimentally verified. *Recall* is defined as TP/(TP+FN), that is, the fraction of EC numbers in EcoCyc that were correctly predicted by the annotation tools. Higher recall values indicate a more complete annotation covering more of the EcoCyc annotations, and lower values indicate EcoCyc EC annotations that were not predicted by the four tools. Thus, a more lenient annotation policy (e.g. merging annotations from all tools) will tend to generate fewer false negatives but more false positives, achieving a higher recall at the expense of lower precision. Conversely, a more restrictive annotation policy (e.g. only including EC numbers if all tools agree on them) can increase precision, but at the expense of a lower recall.

**Figure 4.**
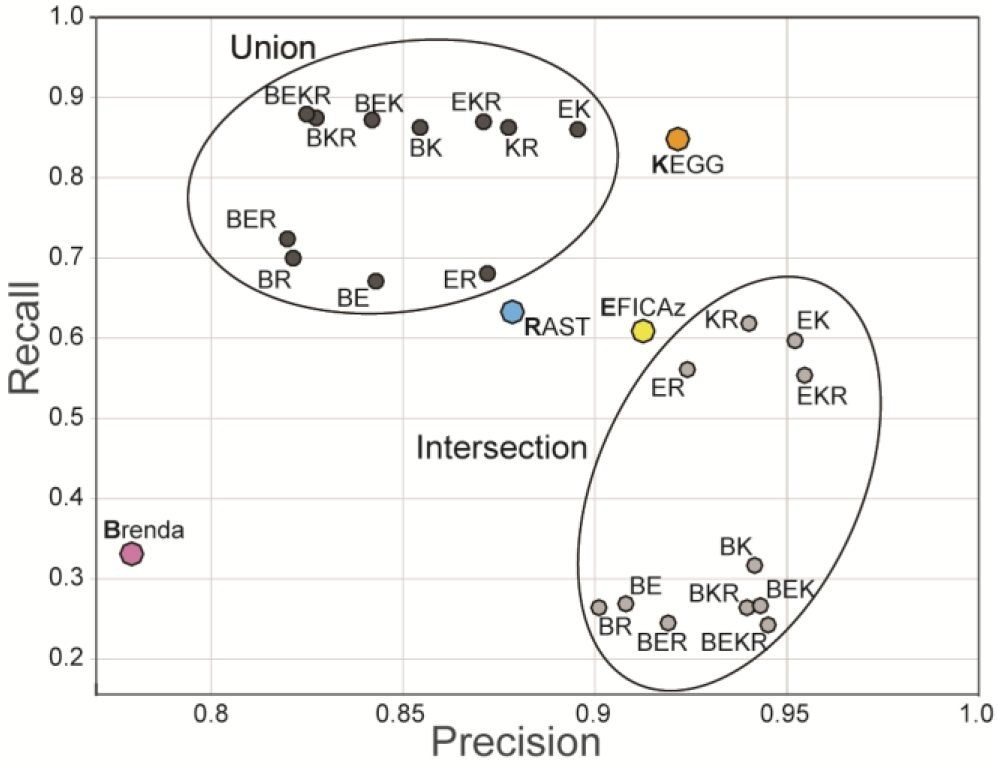
Precision vs Recall of EC numbers for different combinations of tools on EcoCyc. Individual tools are denoted by B, E, K, or R for BRENDA, EFICAz, KEGG, and RAST, respectively.

Figure 4 shows a plot of precision versus recall for all the different combinations of tools. Out of the individual tools, KEGG performed best in terms of both precision and recall on this dataset (although as mentioned before, performance on E. coli K12 may not reflect performance on other genomes), and a simple Blast against the BRENDA database performed worst. Combinations that contain some union of the tools have a higher recall than each of the individual tools in the combination, but a somewhat lower precision (>80%). In contrast, intersections of annotations from two or more tools show very high (>90%) precision but much lower recall (<65%). A consensus annotation that produces both higher recall and higher precision might be achieved by means of a weighted sum of all the annotation sources, similar to the approach taken by EnzymeDetector (33).

Even though we expected the *E. coli* K-12 genome to be a best-case annotation candidate, there were still significant differences in the annotations produced by the different tools, with each tool only covering a subset of the known enzymes in EcoCyc. The four annotation tools annotated a significantly larger fraction of the genome, and showed much more agreement on *E. coli* than on more remote lineages such as Actinomycetes, Bacteroidetes, or Clostridia (Figure 5A). For *E. coli* K-12, 60% of EC numbers were agreed on by 3 or more tools, while 28% EC numbers come from only a single tool. In contrast, for *P. difficile* 630, only 33% of EC numbers were agreed on by 3 or more tools, and 48% of EC numbers come from only a single tool. Compared to the five *E. coli* strains in our dataset, the annotation tools also cover on average around 30% fewer genes for the 13 genomes in the bottom half of Figure 5B. We see a similar effect when we compare the annotation coverage for *B. subtilis* – arguably the best studied Gram-positive model organism – with all 8 other Gram-positive genomes in our dataset. These results suggest that genome coverage for each tool, and agreement in annotations across tools are significantly worse for organisms that are more phylogenetically distant from well-studied model organisms, making it all the more important to combine multiple tools when annotating these genomes.

**Figure 5.**
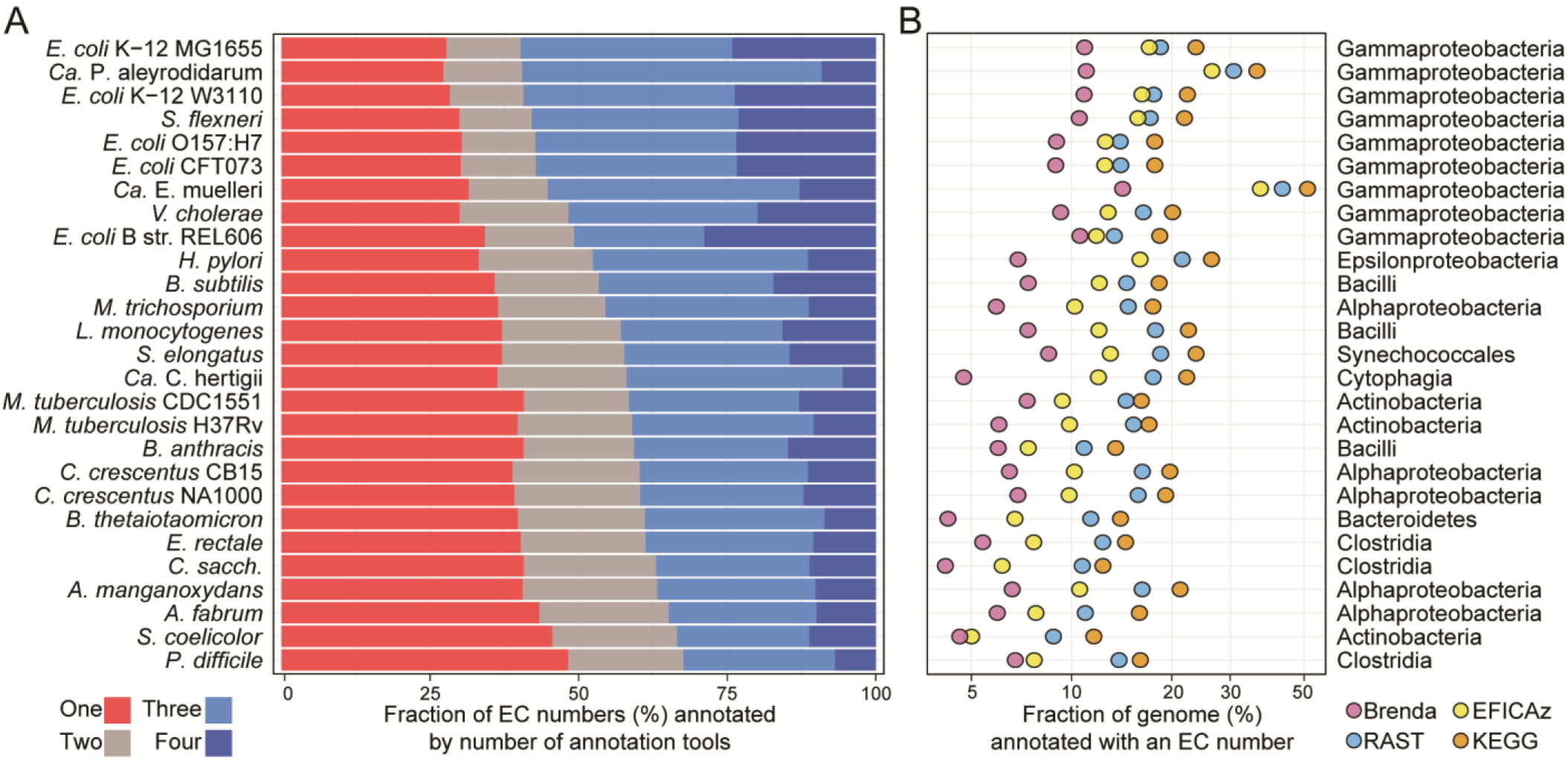
A: Fraction of EC numbers for each genome on which one, two three or all four tools agree. The 27 reference genomes were sorted with respect to the fraction of EC numbers that were predicted by 3 or more tools (blue bars). The top of the list is dominated by model organisms such as *E. coli, B. subtilis*, and closely related organisms. As we move farther away from such well-studied model organisms, the fraction of unique EC numbers predicted only by a single tool (red bars) increases, at the expense of those predicted by multiple tools. B: The fraction of genes annotated as enzymes by each tool likewise decreases as we move farther away from model organisms such as E. coli. Note that two of the organisms with a drastically reduced genome content, *Candidatus* Portiera aleyrodidarum BT-QVLC and *Candidatus* Evansia muelleri, also have a relatively higher fraction of core metabolic enzymes.

### Transporter Annotations

Knowledge of the molecules and substrates an organism can transport and exchange with the environment can help to build a more accurate metabolic model. Both RAST and KEGG include membrane transport annotations, yet both tools yielded on average only 114 and 204 transporter predictions per genome, respectively (Figure 6A and Supplementary Figure S2). Many of these annotated transporters lack substrate predictions (52% of transporter annotations in RAST, 25% in KEGG) or have ambiguous substrate predictions (ranks 3-4 (Table 2); 20% in RAST, 28% in KEGG), while less than half have substrate predictions that are sufficiently detailed to be incorporated in a metabolic model (ranks 1-2; 28% in RAST, 48% in KEGG; Figure 6B). In contrast, TransportDB produces an average of 426 transport annotations per genome, and most of those have specific substrate predictions (59% rank 1-2; 32% rank 3-4, 10% rank 5; Figure 6B).

**Figure 6.**
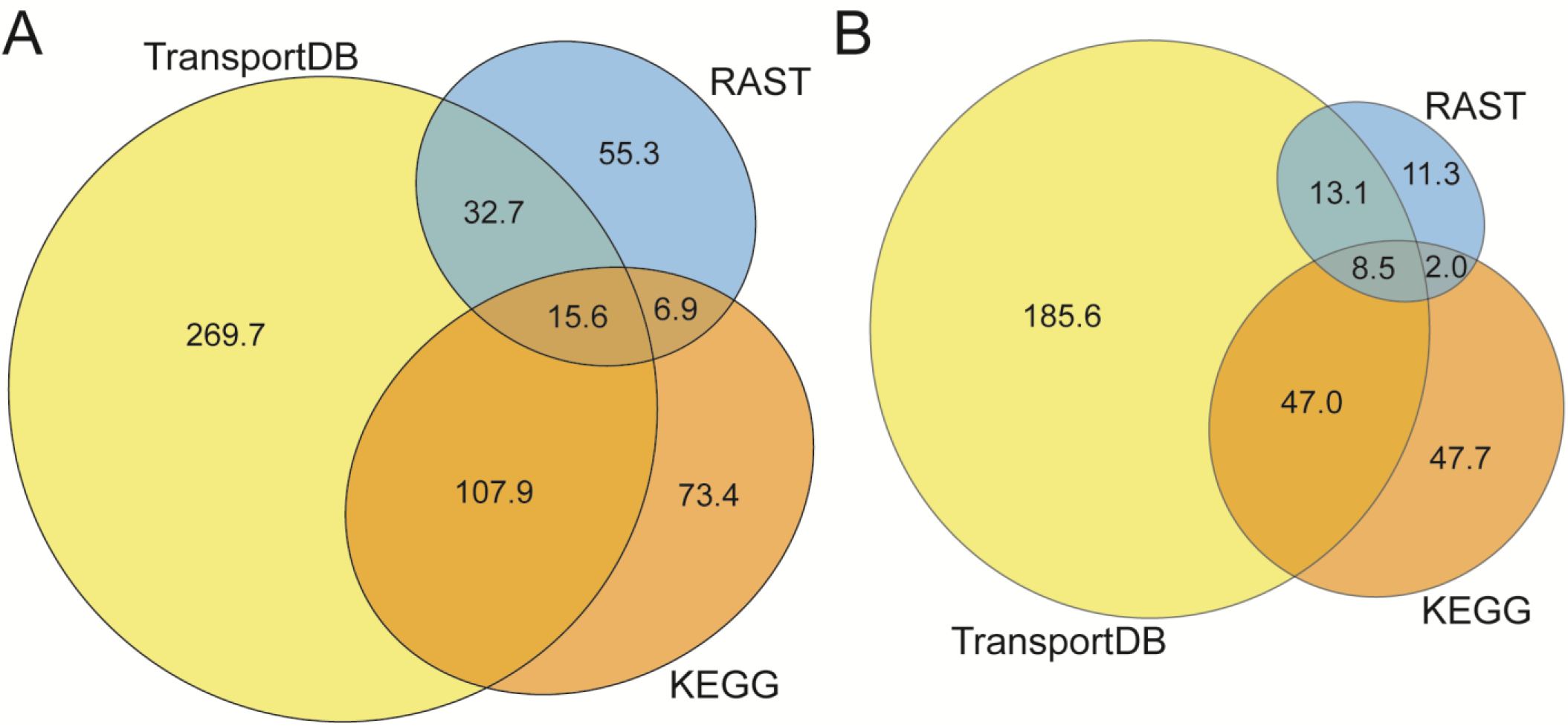
A: Total number of genes annotated as transporters, regardless of substrate. B: Transporter annotations with substrates predictions specific enough to be included in metabolic models (rank 1 or 2).

Transporter annotations by RAST, KEGG and TransportDB showed surprisingly little overlap. Out of the more than 15,000 genes annotated as transporters (regardless of substrate prediction), the three tools only agree on 2.8% (423/15,161). Out of those, only 130 genes are annotated by all three tools with a specific substrate prediction (ranks 1-2). When two or more tools provide a sufficiently specific substrate annotation, the substrate annotations tend to agree 85% of the time, even if they may not be perfectly identical (for example, one transporter was annotated as “leucine/valine”, “leucine”, and “branched-chain amino acid” by TransportDB, RAST and KEGG respectively). Overall, the detailed transporter annotations by TransportDB’s Transporter Automatic Annotation Pipeline provide a significant advance over more general metabolic annotation tools such as RAST and KEGG.

## Conclusions

This analysis has led us to make recommendations for providing a more comprehensive metabolic genome annotation. We found that a single annotation tool is often insufficient unless one is only interested in core metabolism where different tools often agree. Organisms that are phylogenetically farther removed from well-studied model organisms are particularly susceptible, in which case annotation tools will tend to diverge far more. In addition, one can trade off confidence in predictions versus greater coverage by using the intersection or union of multiple annotation tools. BLASTing against a database of reference sequences is generally an inefficient method for annotating enzymes but may be useful to cover more recently assigned EC numbers not yet included by other tools. Still, all these efforts require manual effort to bring together annotation from multiple sources. More tool development is needed to merge annotations beyond simple EC numbers, and a universal reference database for well-balanced reactions and metabolites would be a very valuable resource to merge annotations that use different reaction nomenclatures (42–44). Likewise, now that annotation tools such as TransportDB are producing significant numbers of transporter annotations with substrate predictions that are precise enough to be included in metabolic modeling, more tool development may be needed to fully take advantage of these substrate predictions in Flux Balance Analysis methods, and move beyond the current implicit assumption used by most algorithms that all metabolites can be transported when needed.

## SUPPLEMENTARY DATA

**Supplementary Figure S1.**
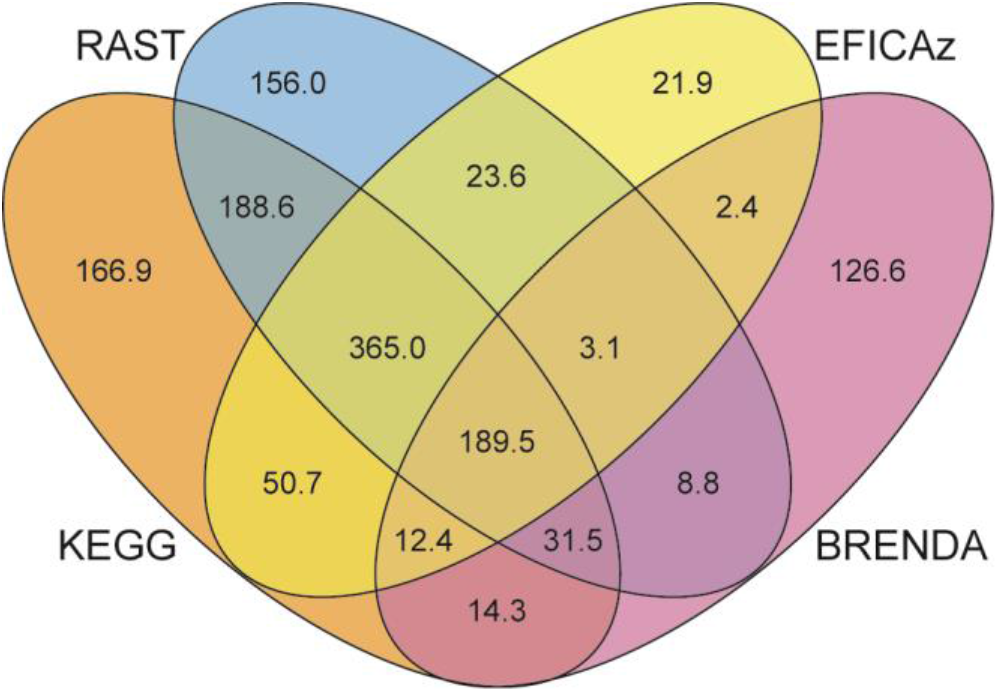
Overlap of annotated genes between the tools (average numbers of genes annotated per genome).

**Supplementary Figure S2.**
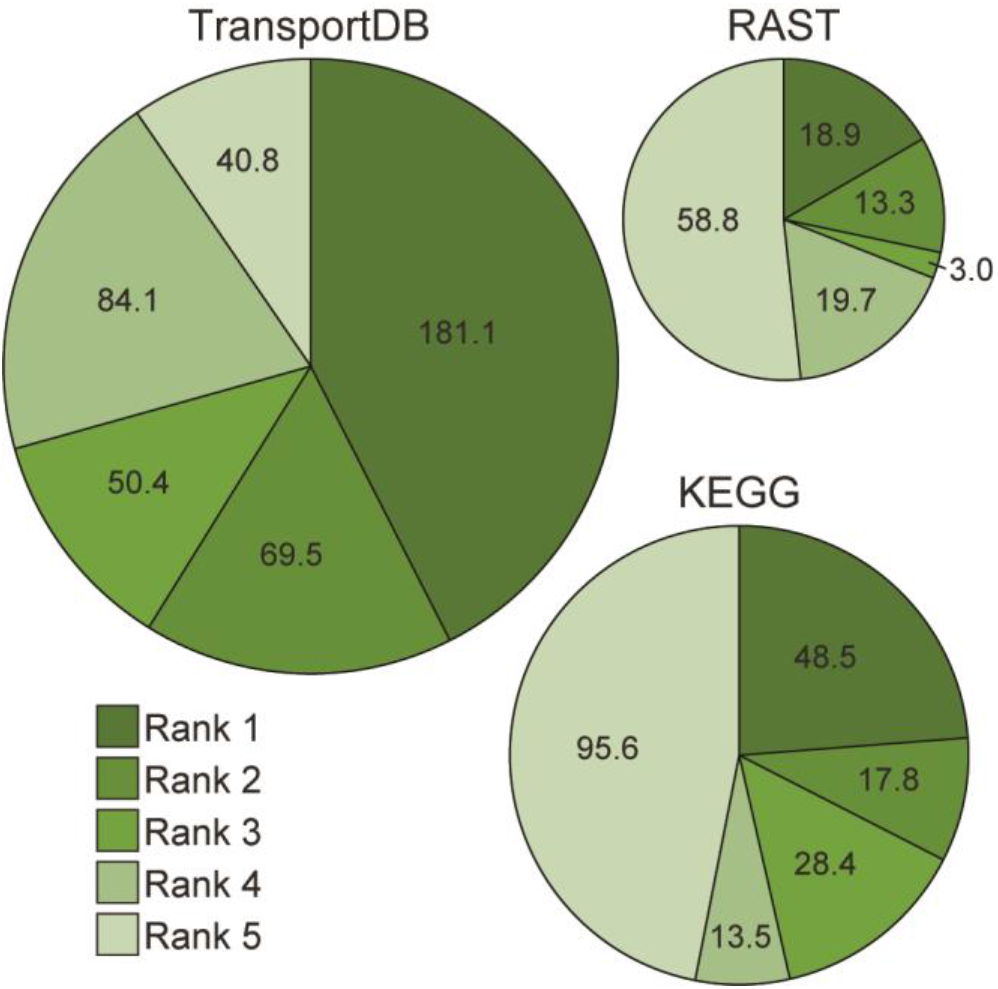
Average transporter annotations per genome produced by TransportDB (426.0), KEGG (203.8) and RAST (113.7) and the distributions of their substrate specificities (rank 1 is most specific, rank 5 has no substrate prediction).

Supplementary Data S3. Zip file with the 27 reference genomes in Genbank format, cleaned to include only coding sequences and locus tags.

Supplementary Data S4. Excel file with all EC annotations by all 4 tools for all 27 genomes.

Supplementary Data S5: Excel file with the EC numbers predicted by all 4 tools in most genomes.

Supplementary Data S6: Excel file with all transporter annotations by all 3 tools for all 27 genomes.

Supplementary Data S7: Excel file with all substrate ranks.

